# Kinase-Inhibitor Binding Affinity Prediction with Pretrained Graph Encoder and Language Model

**DOI:** 10.1101/2025.02.06.636965

**Authors:** Xudong Guo, Zixu Ran, Fuyi Li

## Abstract

**Motivation:** The accurate prediction of inhibitor-kinase binding affinity is crucial in drug discovery and medical applications, especially in the treatment of diseases such as cancer. Existing methods for predicting inhibitor-kinase affinity still face challenges including insufficient data expression, limited feature extraction, and low performance. Despite the progress made through artificial intelligence (AI) methods, especially deep learning technology, many current methods fail to capture the intricate interactions between kinases and inhibitors. Therefore, it is necessary to develop more advanced methods to solve the existing problems in inhibitor-kinase binding prediction.

**Results:** This study proposed Kinhibit, a novel framework for inhibitor-kinase binding affinity predictor. Kinhibit integrates self-supervised pre-trained molecular encoders and protein language models (ESM-S) to extract features effectively. Kinhibit also employed a feature fusion approach to optimize the fusion of inhibitor and kinase features. Experimental results demonstrate the superiority of this method, achieving an accuracy of 92.6% in inhibitor prediction tasks of three MAPK signaling pathway kinases: Raf protein kinase (RAF), Mitogen-activated protein kinase kinase (MEK), and Extracellular Signal-Regulated Kinase (ERK). Furthermore, the framework achieves an impressive accuracy of 93.4% on a dataset containing over 200 kinases. This study provides promising and effective tools for drug screening and biological sciences.

## 1. Introduction

Kinases play a vital role in cell regulation, regulating a variety of cellular processes, including physiological functions such as cell cycle, differentiation, and apoptosis, by catalyzing the transfer of phosphate groups (Cohen, 2002; Fischer, 2013; Cohen, 1982). This key role makes them an important target in drug development, especially in the treatment of cancer and inflammatory diseases (Attwood et al., 2021; Grant, 2009; Kontzias et al., 2012; Lahiry et al., 2010). Predicting the binding affinity of inhibitor-kinases is not only the basis for understanding the function of kinases but also an important prerequisite for designing efficient targeted drugs. However, experimental determination of the binding affinity of inhibitor-kinase is often costly and time-consuming. Therefore, developing efficient computational methods to replace traditional experiments has become a key topic in current research.

Among the many kinase families, the mitogen-activated protein kinase (MAPK) signalling pathway has attracted much attention due to its core regulatory role in cell proliferation, survival, and differentiation, and is the focus of inhibitor design and affinity prediction (Yeung et al., 2018; Haas et al., 2021; Yuan et al., 2020). In recent years, researchers have explored a variety of computational methods to predict the binding affinity of MAPK inhibitors, including traditional machine learning methods such as random forests, support vector machines, Bayesian decision theory, and logistic regression(Xue et al., 2006; Fan et al., 2014; Huang et al., 2015; Li et al., 2018), as well as artificial intelligence methods that have emerged in recent years, such as BatchDTA(Luo et al., 2022), KIPP(Wu et al., 2024), and GPT4Kinase(Liu et al., 2024). These methods have improved the accuracy and efficiency of prediction to a certain extent, but there are still problems such as insufficient feature expression and low performance.

To further improve the performance of MAPK inhibitor binding affinity prediction, this study proposed Kinhibit, a new framework combining graph contrastive learning and pre-trained language models. Specifically, we used graph contrastive learning to pre-train the molecular graph encoder used for inhibitor characterization to fully explore the graph structure features of the inhibitor. At the same time, based on the structure-informed protein language model (ESM-S)(Zhang et al., 2024), the features of kinases were extracted to capture their structure and sequence information. Subsequently, the graph features of the inhibitor and the features of the kinase were fused through a feature fusion approach to capture the complex interaction information between the inhibitor and the kinase. In addition, we used a multi-layer perceptron (MLP) to design a classifier and trained a series of regression models based on the machine learning algorithm using the learned features for the classification of kinase inhibitors and inhibitor-kinase binding affinity prediction, respectively.

Overall, this framework not only significantly improved the accuracy of MAPK inhibitor affinity prediction by introducing the structure-informed features of kinases and the graph structure features of molecules to synergistically optimize the model, but also provided a novel framework for studying the identification and design of inhibitors for other kinases. The results of this study will provide important support for the development of kinase-targeted drugs, especially for the treatment of kinase-related diseases such as cancer.

## 2. Material and Methods

### 2.1. Dataset

In this study, we used the kinase dataset from GPT4Kinase(Liu et al., 2024), which includes information such as the binding domains’ sequences of kinases, the inhibitor SMILES strings, and Kd values of their corresponding inhibitors from the BindingDB database(Gilson et al., 2016). These datasets include the MAPK dataset and the multiple kinase dataset. The MAPK dataset includes three kinases: RAF (EC EC2.7.11.1) includes RAF protooncogene serine/threonine-protein kinase and serine/threonine-protein kinase B-raf, MEK (EC EC2.7.12.2) includes dual specificity mitogen-activated protein kinase kinase 1 and dual specificity mitogen-activated protein kinase kinase 2, ERK (EC EC2.7.11.24) includes mitogen-activated protein kinase 3 and mitogen-activated protein kinase 1 in the MAPK signaling pathway. The MAPK dataset used to train the model is obtained from the initial dataset by removing samples with uncertain Kd values, SMILES duplicates and balancing the labels, including 487 inhibitors: 161 for RAF kinases, 131 for MEK kinases, and 195 for ERK kinases. Similar to the clean process of the MAPK dataset, the Multiple Kinase dataset includes 318 kinases and 3234 inhibitors. To evaluate the model’s prediction effect objectively and intuitively, these datasets were labeled using two strategies based on the distribution range of the Kd values, and divided into a training set and a validation set in a ratio of 8:2(Liu et al., 2024). More details of labeling strategies are shown in Supplementary Table 1.

### 2.2. Overview of the methods

Kinhibit consists of two main processes: Pre-training and finetuning, as depicted in Fig1. The Pre-training phase aims to train a robust small molecular encoder using a graph contrastive learning strategy. The initial input for this step is the multiview SMILES strings (the different SMILES strings of ligand) of the ligands, which are subsequently converted to the molecular graph representation using the RDKit tool(Landrum, 2013) and used as input to the molecular encoder to learn the highdimensional representation of the ligand. Finally, the molecular encoder is optimized by minimizing the contrastive loss between the high-dimensional representation of the ligand. In this study, the molecular encoder used is the E(n) Equivariant Graph Neural Network (EGNN)(Satorras et al., 2021) proposed by Satorras et al.

**Fig. 1.**
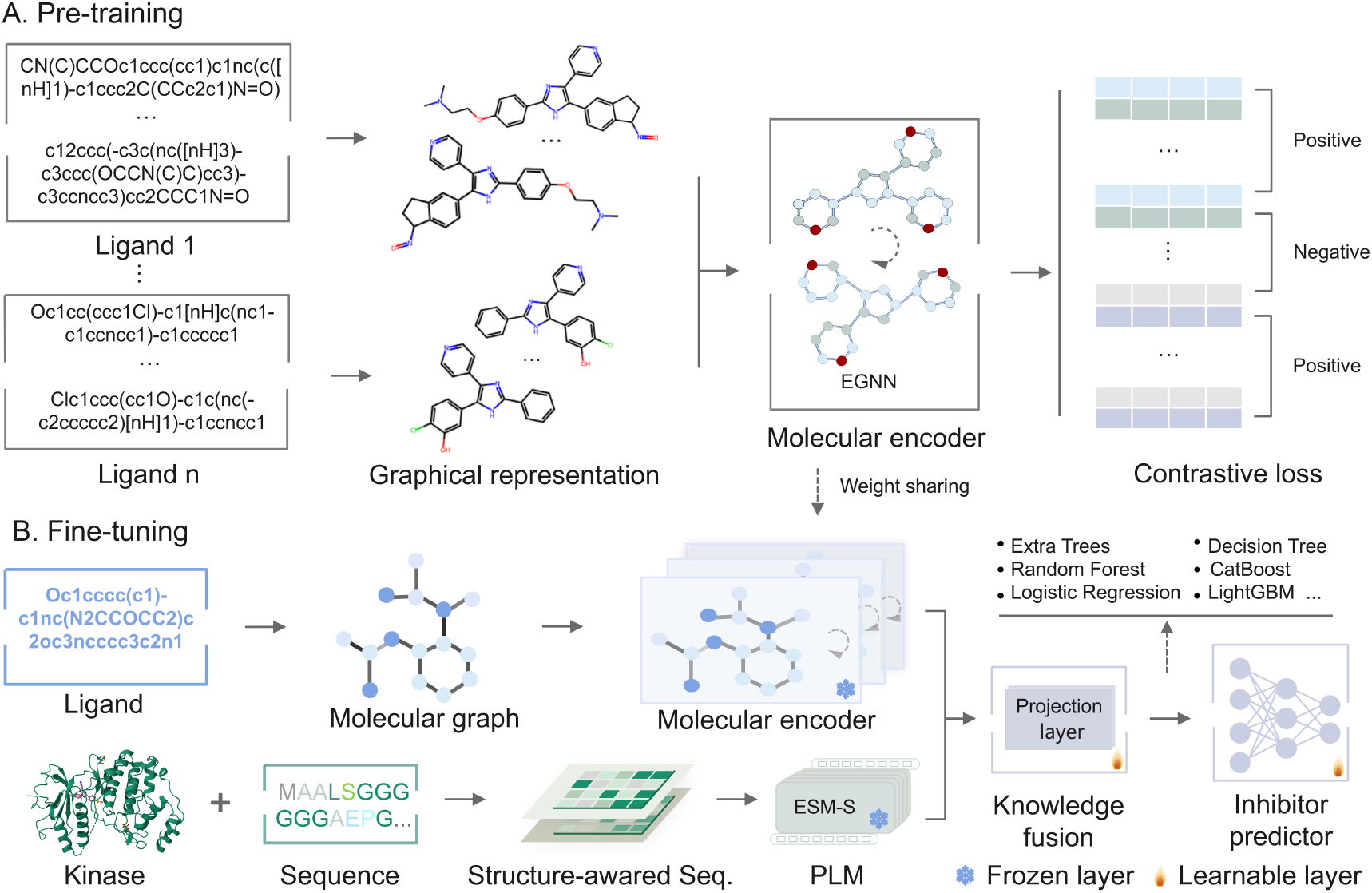
The framework of Kinhibit. A.The process of self-supervised contrastive learning of molecular encoder. B. The process of supervised fine-tuning.

Building on the pre-trained molecular encoder, we integrated the structure-informed protein language model ESM-S to capture high-dimensional representations of kinases. To achieve effective knowledge fusion, projection layers were introduced(Du et al., 2024), which take the high-dimensional representations of kinases and inhibitors as input. This process enables the projection layers to learn efficient ligand representations that are enriched with kinase-specific information. These enriched ligand representations will then be used to train robust inhibitor identification models. In this study, we employed a fine-tuning strategy to achieve this goal. During the fine-tuning phase, the weights of the molecular encoder and ESM-S-based encoder are frozen to retain their pre-trained representations, and the projection layers for both encoders (molecular encoder and ESM-S) to refine and align the embeddings of ligands and kinases in the same high-dimensional space. An end-to-end deep learning approach is employed to update the weights of the projection layers and the inhibitor prediction module. This strategy enables the training of a robust inhibitor classification model, leveraging the advantages of pre-trained features while optimizing task-specific performance. Additionally, the features learned by the model are utilized to train a kinase-inhibitor binding affinity predictor using machine learning algorithms.

### 2.3. Representation of kinases and ligands

The protein language model (PLM) is widely used in the field of bioinformatics due to its outstanding performance in protein representation. ESM is a PLM project initiated by Meta Fundamental AI Research (FAIR) in 2019(Rao et al., 2019). Based on ESM, Tang et al. introduced the integration of remote homology detection to extract structural information into the protein language model and proposed a structure-informed protein language model ESM-S. Like ESM-2, this model supports protein feature extraction in multiple dimensions (320, 480, 640,1280)(Zhang et al., 2024). In this study, the kinases were represented numerically using ESM-S-1280 models, i.e., the 1280-dimensional vectors. For each ligand, we utilize RDKit to convert the SMILES strings into a graph representation. Specifically, the molecular graph of ligand is defined as *G* = (*V, E*), where *V* denotes node sets, each node represents an atom; *E* denotes the edges set, and each edge represents the chemical bonds between atoms. In this study, each node of the molecular graph *G* is characterized by a feature vector containing 8 attributes, including atom type, degree, formal charge, chirality, number of *H*, hybridization, aromaticity, and atomic mass. Similarly, each edge is characterized by four types of attributes, including bond type, ring, stereo, and conjugated.

### 2.4. Pre-training and fine-tuning

In this study, we utilized self-supervised contrastive learning to enhance the molecular encoder’s ability to distinguish between different ligand’ SMILES strings. For each ligand, we applied the RDKit tool to generate augmented graphs of ligands by performing the following operations: generating multi-view SMILES strings of ligands (augmented ligand’s SMILES) and converting the SMILES to molecular graphs. The augmented graphs for the same ligand are treated as positive pairs, while other ligands’ SMILES in the batch data are considered negative pairs. With these augmented views, we pre-train the molecular encoder using the self-supervised contrastive learning loss function *L*_*self*_ (Khosla et al., 2020; Jaiswal et al., 2020). During training, we applied RDKit to calculate the similarity between samples in the batch data using the SMILES strings of the ligands. The pseudocode for self-supervised pre-training is shown in Algorithm 1. The *N* denotes the number of original ligand’s SMILES and 10*N* denotes the number of augmented SMILES.

#### Algorithm 1 Pseudocode of molecular encoder pre-training

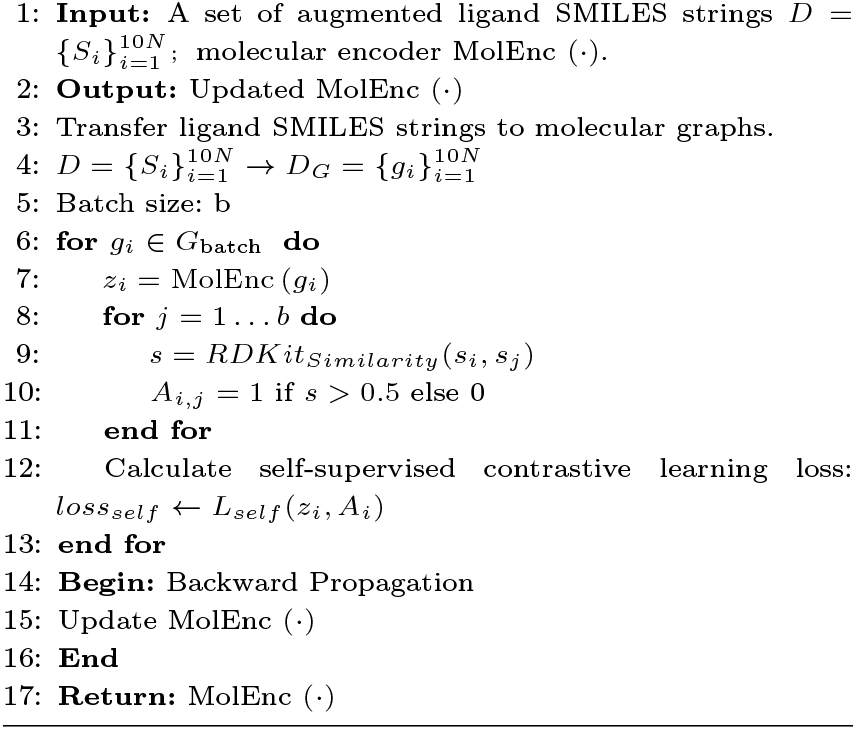

In the fine-tuning phase, during forward propagation, the ligand’s SMILES and the kinase’s sequence and position information are fed into the pre-trained molecular encoder and structure-informed protein language model ESM-S, respectively, to generate embeddings. These embeddings are then passed into the projection layers, which act as the knowledge fusion module to learn an efficient representation of ligands. These ligand’s representations are then used as inputs to an inhibitor classifier consisting of multiple fully connected layers to generate predictive values. During backpropagation, the classification cross-entropy was used as the loss function to calculate the loss values of predicted and true labels. We employ the Adam optimizer with a learning rate of 0.001 to minimize the distance between each class label and the predicted probability value. It is to be noted that, in this stage, the molecular encoder and ESM-S are frozen so that it does not update the learned weights, and only the weights of the projection layers and the inhibitor classifier will be updated. The loss function is shown below:

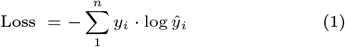

Where *y*_*i*_ takes the value 0, 1, or 2, and *ŷ*_*i*_ takes the value in [0,1]. When the *y*_*i*_ value is 0, it means that the ligand sample *s*_*i*_ with low affinity; on the contrary, if the *y*_*i*_ value is 1, it means that the ligand sample *s*_*i*_ with medium affinity; if the y value is 2, it means that the ligand sample *s*_*i*_ with high affinity. *n* is the number of classes (*n* = 3). In this study, to prevent overfitting, we use dropout as well as set the number of EGNN layers to 1. The input dimension, hidden layer dimension, and output dimension of the EGNN are set to 133. The two input dimensions of the projection layer are 1,280 and 133, respectively, and the output dimension is 128. The input dimension of the fully connected layer is 256, and the output dimension is 3, which is equivalent to the number of label types.

## 3. Results and discussion

### 3.1. Kinhibit’s multi-class performance

In this study, we adopted the labeling strategy proposed in the literature(Liu et al., 2024). Table 1 shows the classification performance comparison between Kinhibit and GPT4Kinase under the two labeling strategies. As shown in Table 1, Kinhibit outperformed GPT4Kinase in accuracy, precision, and F1 score across the three datasets: ERK, MEK, and RAF. Specifically, under labeling strategy A, Kinhibit achieved an accuracy of 0.912 on the RAF dataset, with precision and F1 scores of 0.886 and 0.899, respectively, surpassing GPT4Kinase’s scores of 0.627 and 0.480. On the ERK dataset, Kinhibit achieved accuracy, precision, and F1 scores of 0.895, 0.891, and 0.889, while GPT4Kinase scored 0.744, 0.819, and 0.758, respectively. On the MEK dataset, Kinhibit recorded scores of 0.889, 0.887, and 0.886, while GPT4Kinase’s scores were 0.815, 0.600, and 0.814. Under the labeling strategy B, Kinhibit and GPT4Kinase performed relatively smoothly in accuracy,prcision, and F1 score, but Kinhibit outperformed GPT4Kinase in accuracy by nearly 24 percentage points. In terms of AUC, under labeling strategy A, Kinhibit achieved the highest AUC value of 0.956 on the RAF dataset, surpassing GPT4Kinase by approximately 35 percentage points. Under labeling strategy B, Kinhibit reached a maximum AUC value of 0.9 on the ERK dataset, which is nearly 10 percentage points higher than GPT4Kinase. Overall, Kinhibit’s performance on the three datasets is relatively stable under the two labeling strategies, while GPT4Kinase’s performance is relatively stable under the labeling strategy B.

**Table 1.** Comparison of classification performance with the GPT4Kinase model under different labeling strategies.

| Kinase | Method | Accuracy |  | Precision |  | F1 Score |  | AUC |  |
| --- | --- | --- | --- | --- | --- | --- | --- | --- | --- |
|  |  | Strategy-A | Strategy-B | Strategy-A | Strategy-B | Strategy-A | Strategy-B | Strategy-A | Strategy-B |
| ERK | Kinhibit | 0.895 | 0.868 | 0.891 | 0.877 | 0.889 | 0.870 | 0.895 | 0.900 |
|  | GPT4Kinase | 0.744 | 0.692 | 0.819 | 0.788 | 0.758 | 0.719 | 0.852 | 0.810 |
| MEK | Kinhibit | 0.889 | 0.778 | 0.887 | 0.805 | 0.886 | 0.772 | 0.954 | 0.817 |
|  | GPT4Kinase | 0.815 | 0.741 | 0.600 | 0.765 | 0.814 | 0.738 | 0.848 | 0.782 |
| RAF | Kinhibit | 0.912 | 0.882 | 0.886 | 0.885 | 0.899 | 0.876 | 0.956 | 0.882 |
|  | GPT4Kinase | 0.888 | 0.873 | 0.627 | 0.881 | 0.480 | 0.846 | 0.609 | 0.676 |

### 3.2. Performance of affinity prediction based on Kinhibit’s features

As can be seen from the above, Kinhibit shows stable performance under both labeling strategies, which may be derived from its strong feature learning ability. Therefore, we used the learned features and the PyCaret tool(Olanipekun and Mashao, 2024) to further evaluate the affinity prediction performance of the model. Specifically, we first saved the input of the inhibitor classifier of Kinhibit and obtained the embeddings of the four data sets, including training sets and test sets. Then, we used the PyCaret tool to compare multiple regression models on the training set to obtain the best regression models and used the Kd values conversion standard given in the literature(Liu et al., 2024) to convert the predicted values, that is, normalize the predicted Kd values, mark 0 =*<* Normalized value *<*0.3 as 0, indicating low affinity, mark 0.3=*<* Normalized value*<*0.7 as 1, indicating medium affinity, and mark 0.7=*<* Normalized value*<*1 as 2, indicating high affinity. Finally, the regression model was evaluated using accuracy, precision, and F1 Score.

As shown in Table 2, we compared Kinhibit with four other methods: GPT4Kinase, AutoDock(Trott and Olson, 2010), BatchDTA, and KIPP. Specifically, under labeling strategy B, we evaluated the trained optimal regression model (the best regression model for each dataset) against these four tools, utilizing the above Kd value conversion standard. The results in Table 2 indicate that Kinhibit achieves superior overall affinity prediction performance compared to the other methods. On the ERK dataset, Kinhibit achieves the highest accuracy and F1 Score, outperforming all competitors. GPT4Kinase ranks fourth on the accuracy, trailing behind BatchDTA and KIPP. In terms of precision, Kinhibit ranks second (0.753), slightly below GPT4Kinase (0.788) but higher than BatchDTA, KIPP, and AutoDock. On the MEK dataset, Kinhibit outperforms all other methods across all three metrics, with scores that are 17 percentage points higher than GPT4Kinase, which ranks second. For the RAF dataset, Kinhibit achieves accuracy and F1 Scores of 0.853 and 0.846, respectively, which are slightly lower than GPT4Kinase’s 0.873 and 0.846, placing Kinhibit in second. However, in terms of precision, Kinhibit achieves a higher value (0.890) compared to GPT4Kinase (0.881). Finally, on the Multi-Kinase dataset, Kinhibit outperforms GPT4Kinase across all three metrics, with an average improvement of 21 percentage points.

**Table 2.** Comparison results of evaluation metrics of Kinhibit, GPT4Kinase, AutoDock Vina, BatchDTA, and KIPP in the prediction of inhibitor-kinase affinity.

| Metrics | Method | ERK | MEK | RAF | Multi-Kinase |
| --- | --- | --- | --- | --- | --- |
| Accuracy | Kinhibit | **0.789** | **0.926** | 0.853 | **0.854** |
|  | GPT4Kinase | 0.692 | 0.741 | **0.873** | 0.662 |
|  | AutoDock | 0.211 | 0.259 | 0.177 | - |
|  | BatchDTA | 0.718 | 0.519 | 0.294 | - |
|  | KIPP | 0.711 | 0.519 | 0.529 | - |
| Precision | Kinhibit | 0.753 | **0.926** | **0.890** | **0.934** |
|  | GPT4Kinase | **0.788** | 0.765 | 0.881 | 0.729 |
|  | AutoDock | 0.244 | 0.260 | 0.108 | - |
|  | BatchDTA | 0.562 | 0.333 | 0.451 | - |
|  | KIPP | 0.586 | 0.431 | 0.424 | - |
| F1 Score | Kinhibit | **0.765** | **0.926** | 0.842 | **0.892** |
|  | GPT4Kinase | 0.719 | 0.738 | **0.846** | 0.656 |
|  | AutoDock | 0.162 | 0.254 | 0.148 | - |
|  | BatchDTA | 0.579 | 0.365 | 0.313 | - |
|  | KIPP | 0.607 | 0.467 | 0.461 | - |

### 3.3. Comparison of Molecular Graph Features and Knowledge Fusion Features

To evaluate the effectiveness of the feature representations learned by Kinhibit for affinity prediction, we analyzed three types of ligand features on the Multi-kinase dataset: molecular graph features, molecular fingerprint features, and knowledge fusion features. The molecular graph features are extracted from the output of the self-supervised pre-trained molecular encoder, molecular fingerprint features are extracted from smiles using RDKit, and the knowledge fusion features are extracted from the output of the projection layers. The results are illustrated in Fig 2, where we show the t-SNE(Van der Maaten and Hinton, 2008) visualizations where data points are colored by affinity level (low, medium, and high). The clusters shown in Fig 2A show relatively weak separation, indicating that these self-supervised learned features have limited discriminative power in distinguishing affinity levels. Similarly, the molecular fingerprint features have limited discriminative power for some ligands (Fig 2B). Fig 2C shows the t-SNE visualization of the ligand features fused with kinase information. Compared with the others, the clusters are more clearly separated, but they also show the same situation as the molecular fingerprint features, that is, the separation is relatively low on some ligands, which may be caused by inaccurate thresholds for distinguishing between high, medium and low affinities. The results show that the knowledge fusion feature can capture the potential patterns related to the affinity level.

**Fig. 2.**
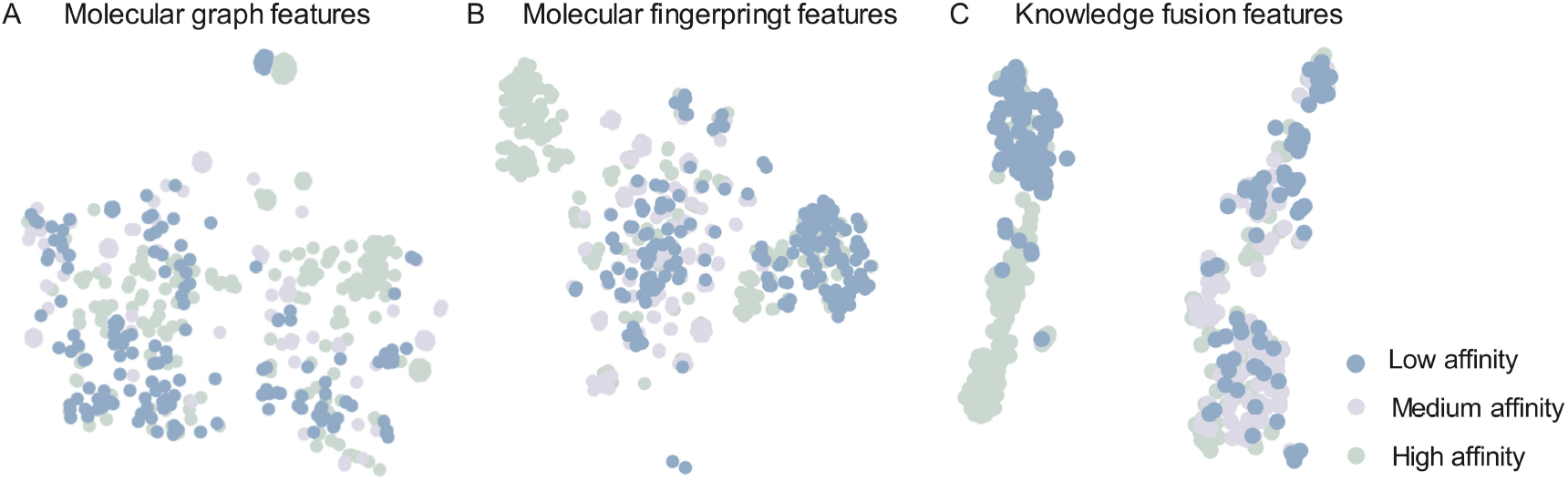
The t-SNE visualization of different features. A. Visualization of molecular graph features. B. Visualization of molecular fingerprint features.

Fig 3A shows the performance of the best regression model trained using two features (molecular graph features and knowledge fusion features) on four key metrics (accuracy, precision, recall, and F1 score), as well as the distribution of predicted and true affinity. The results show that, except for Precision, the regression model using features fused with kinase information is better than the model using molecular graph features in the other three metrics. Similarly, Fig 3B shows that the distribution of binding affinity predicted by the regression model using features fused with kinase information is closer to the true distribution. In summary, these results demonstrate the effectiveness of Kinhibit and suggest that incorporating kinase information into the feature set can improve the performance of kinase inhibitor affinity prediction.

**Fig. 3.**
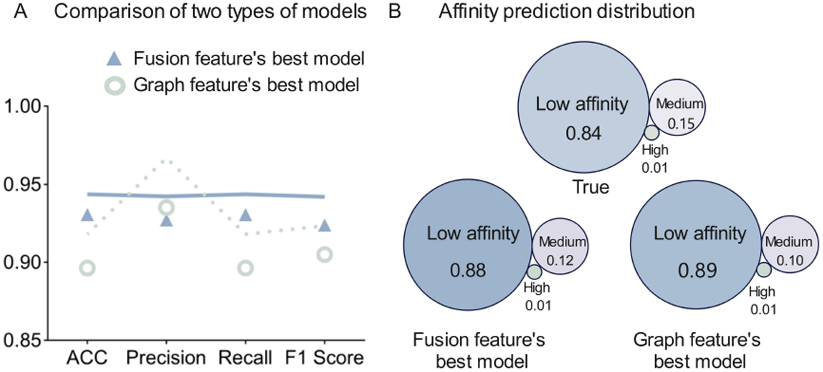
Comparison of regression models trained with two features (molecular graph features and knowledge fusion features). A. Performance comparison on four key metrics. B. Distribution comparison of predicted and true affinities.

### 3.4. Model interpretation

To validate the effectiveness of the multi-view graph comparison learning strategy, we analyze the molecular encoder using the attention mechanism. Specifically, we input the graph representation of the ligand into the molecular encoder and extract the output of the attention layer in the edge feature learning module as a measure of the importance of the edges, which is subsequently ranked. The top 20 edges are considered as the important edges predicted by the Kinhibit. As shown in Table 3, we selected low-, medium-, and high-affinity inhibitors in the test set of three datasets (ERK, MEK, and RAF). According to the labeling strategy B, all these inhibitors were correctly predicted by Kinhibit and the corresponding affinity predictors yielded predictions that were closer to the true Kd values. And, these inhibitors, exhibit a situation consistent with previous studies(Liu et al., 2024), i.e., high-affinity inhibitors tend to have multiple important rings relative to those with lower affinity. These rings contribute to the formation of rigid molecular structures that can stabilize the conformation. As shown in Fig 4, we have highlighted important edges predicted by Kinhibit on the 2D structural graphs of these ligand molecules. As can be seen, most of the important edges predicted by Kinhibit are on the heterocyclic rings of the molecules, and the heterocyclic ring plays an important role in the binding of the inhibitor to the active site of the kinase, which does not only stabilize the molecular conformation but also provides a specific binding site. However, in some medium-affinity inhibitors (e.g., ERK-1 and MEK-1), the significant edges (ring) predicted by Kinhibit are near the end, not the middle heterocyclic rings. This suggests that for Kinhibit, heterocyclic ring information is not an important factor in predicting it as a medium affinity inhibitor. In addition, some of the important edges predicted by Kinhibit, such as ERK-2, MEK-2, and RAF-2, are on bonds containing nitrogen or oxygen atoms that can form hydrogen bonds to increase the affinity of the molecule. To further analyze the interpretability of the model, we calculated and counted the important edges predicted by Kinhibit for all ligands in the test sets of MEK, ERK, and RAF. As shown in Fig. 5A-5C, we show the distribution of important edges predicted by Kinhibit on ligands with different affinities, including AROMATIC edges, SINGLE edges, and DOUBLE edges, as well as the distribution of atom pairs that constitute these edges. From the figure, we can find that on ligands with high and medium affinity, AROMATIC edges account for the majority of the predicted important edges, reaching 62% and 55% respectively, followed by SINGLE, accounting for 35% and 43% respectively, and finally DOUBLE, accounting for 3% and 2% respectively. However, on low-affinity ligands, among the important edges predicted by Kinhibit, SINGLE edges account for 44%, second only to AROMATIC edges at 45%, and DOUBLE edges are significantly improved compared to ligands with high and medium affinity, reaching 11%. From this, we infer that in the ligand molecule, the AROMATIC bond plays an important role in the binding process between the ligand and the kinase, which may be because it is usually distributed on the benzene ring and other conjugated ring systems, and is closely related to the stability of the molecule and the π-π interaction. The DOUBLE bond accounts for a larger proportion in the ligands with low affinity than in the ligands with high and medium affinity, suggesting that it is an important factor affecting the binding of the ligand to the kinase, possibly because it increases the rigidity of the molecule and also affects the conformational changes of the ligand during the binding process with the kinase. In summary, we infer that Kinhibit benefits from the multi-view graph contrastive learning strategy, which can capture important structural information about molecules.

**Table 3.**
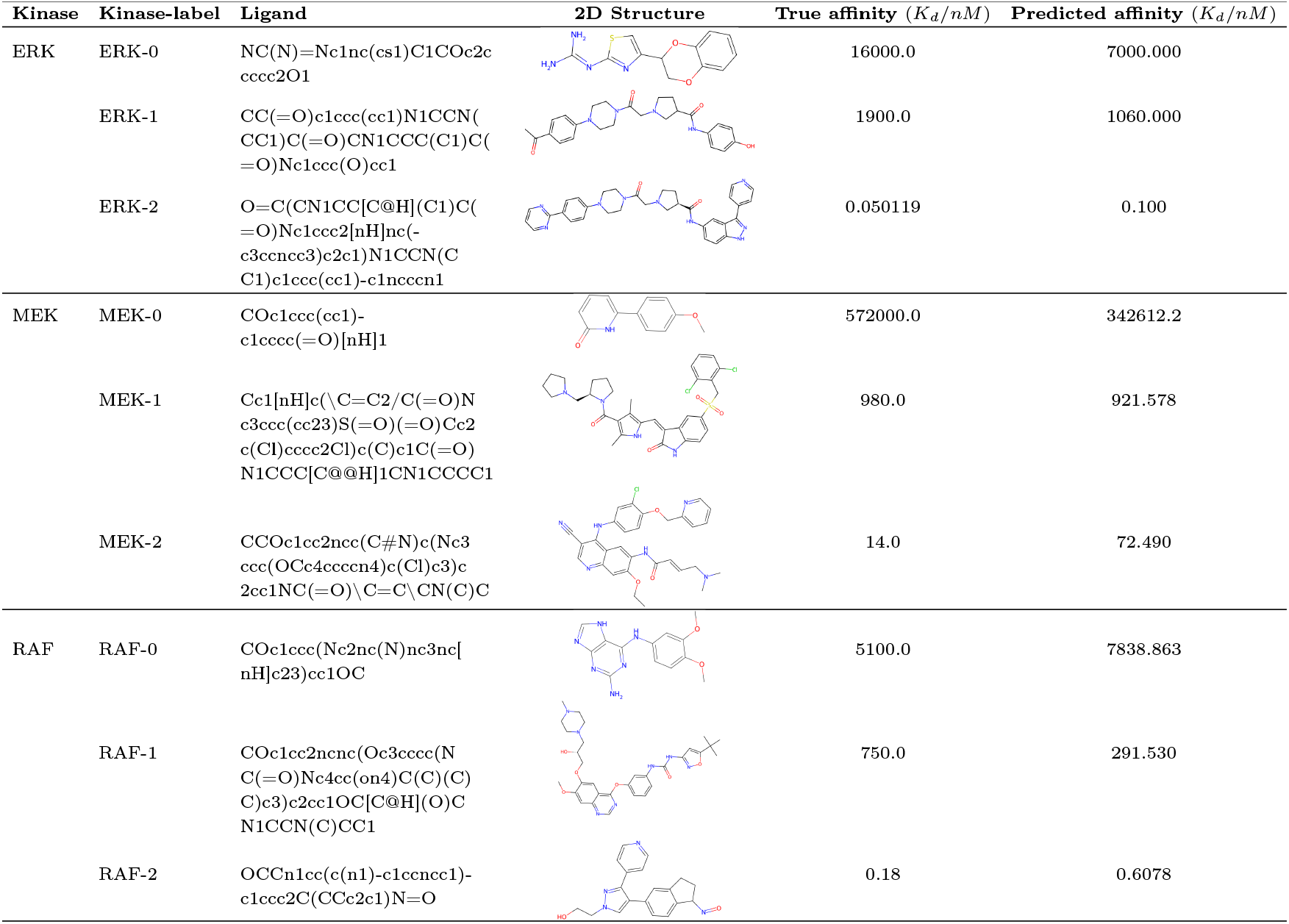
Representative molecular information with different affinities predicted by Kinhibit.

**Fig. 4.**
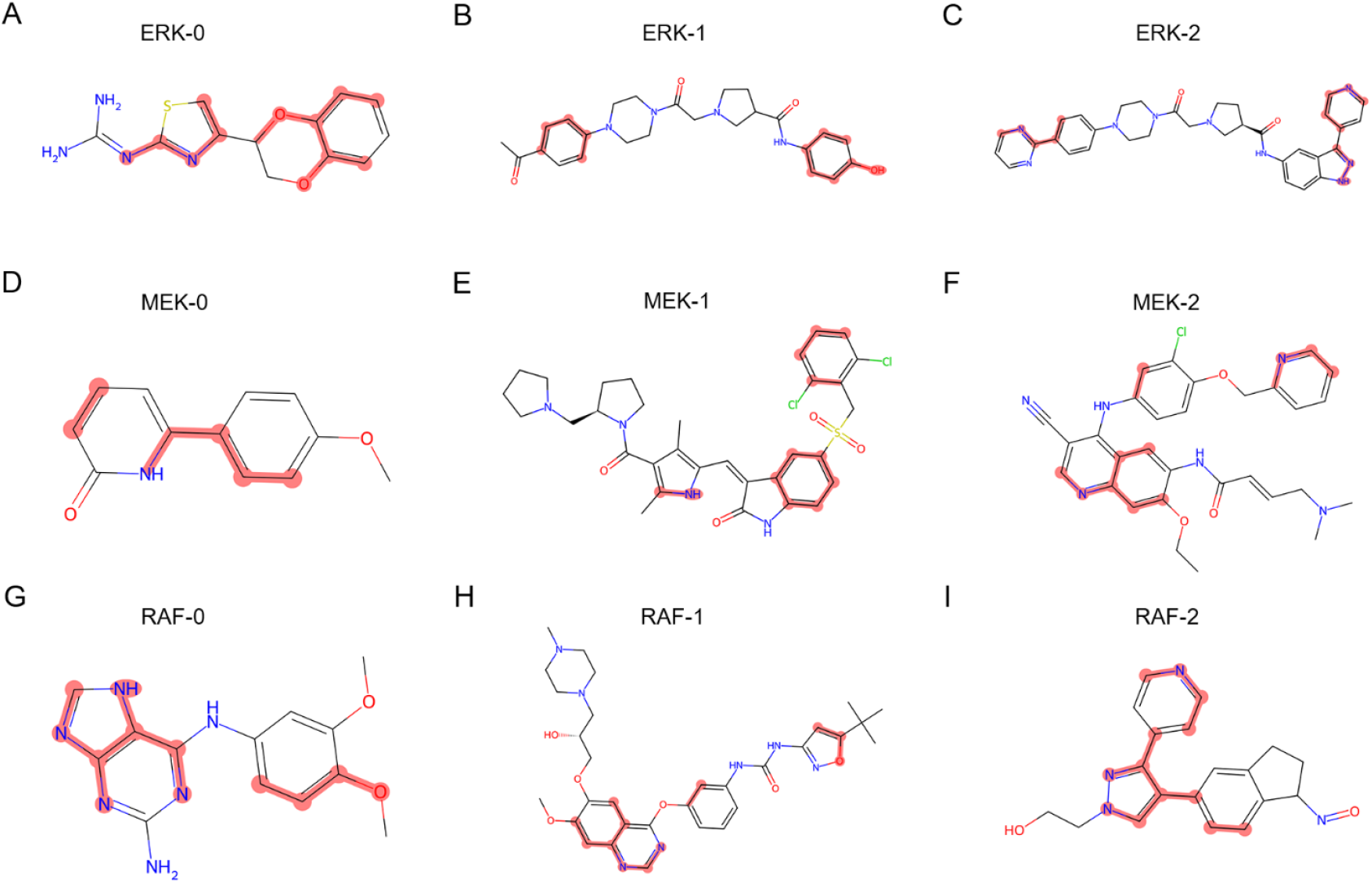
Contribution edges (rings) of inhibitor predicted by Kinhibit. A. Ligands with low- (ERK-0), medium- (ERK-1), and high-affinity (ERK-2) in the ERK test dataset. B. Ligands with low- (MEK-0), medium- (MEK-1), and high-affinity (MEK-2) in the MEK test dataset. A. Ligands with low- (RAF-0), medium- (RAF-1), and high-affinity (RAF-2) in the RAF test dataset.

**Fig. 5.**
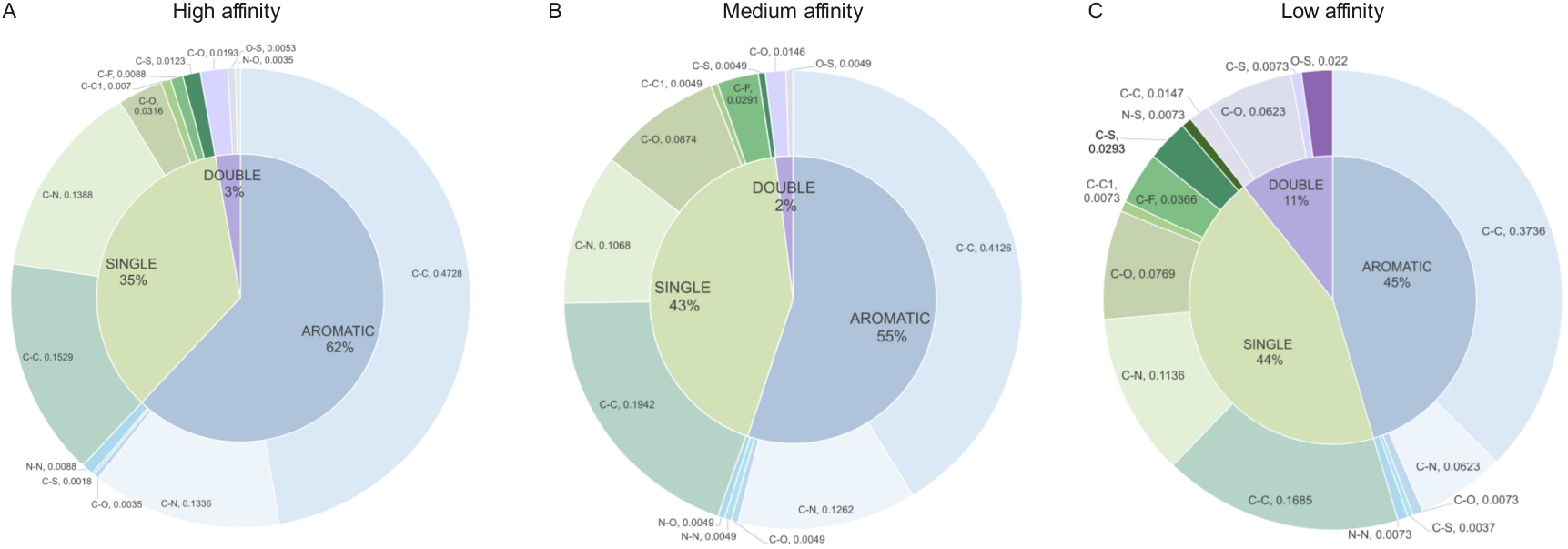
Distribution of important edges predicted by Kinhibit at different affinities.

### 3.5. Case study

To further verify the ability of Kinhibit to identify kinase inhibitors, we used AutoDock for molecular docking. As shown in Fig 6, we selected two examples from the test set for molecular docking verification, namely the docking of ligand-58985117 (PubChem CID: 58985117) with protein Mitogen-activated protein kinase 1 (UniProt accession id: P28482, PDB: 2Y9Q), and the docking of ligand-5330790 (PubChem CID: 5330790) with protein Dual specificity mitogen-activated protein kinase kinase 2 (UniProt accession id: P36507, PDB: 1S9I). The ligand-58985117 exhibits potent inhibitory activity against Mitogen-activated protein kinase 1 (MKPK1), which mediates a variety of biological functions including cell growth, adhesion, survival, and differentiation. Many of its substrates are responsible for processes such as translation, mitosis, and apoptosis(Yoon and Seger, 2006; Yao and Seger, 2009; Wortzel and Seger, 2011). The ligand-5330790 exhibits potent inhibitory activity against cell cycle protein-dependent kinases (CDK1, CDK2), Mitogen-activated protein kinase 1, etc., and inhibits in vitro cell proliferation of various human tumor cells(Lin et al., 2005). As shown in Fig 6A, ligand-58985117 is in the binding pocket formed by amino acids GLU-33, VAL-39, LYS-54, GLN-105, ASP-111, LYS-114, and ASP-167 of protein 2Y9Q and interacts with them to form a hydrogen bond. We found that predicted key edges (rings) or atoms are strongly associated with these formed bonds. Similarly, the docking results of ligand-5330790 with protein 1SPI (Fig 6B) show that the ligand interacts with amino acids ALA-80, GLY-81, GLY-83, HIS-104, and MET-223 and that the predicted critical edges or rings are associated with these interactions. In summary, we infer that the superior performance of Kinhibit over other methods may stem from Kinhibit’s ability to recognize atoms or rings that are strongly associated with interactions.

**Fig. 6.**
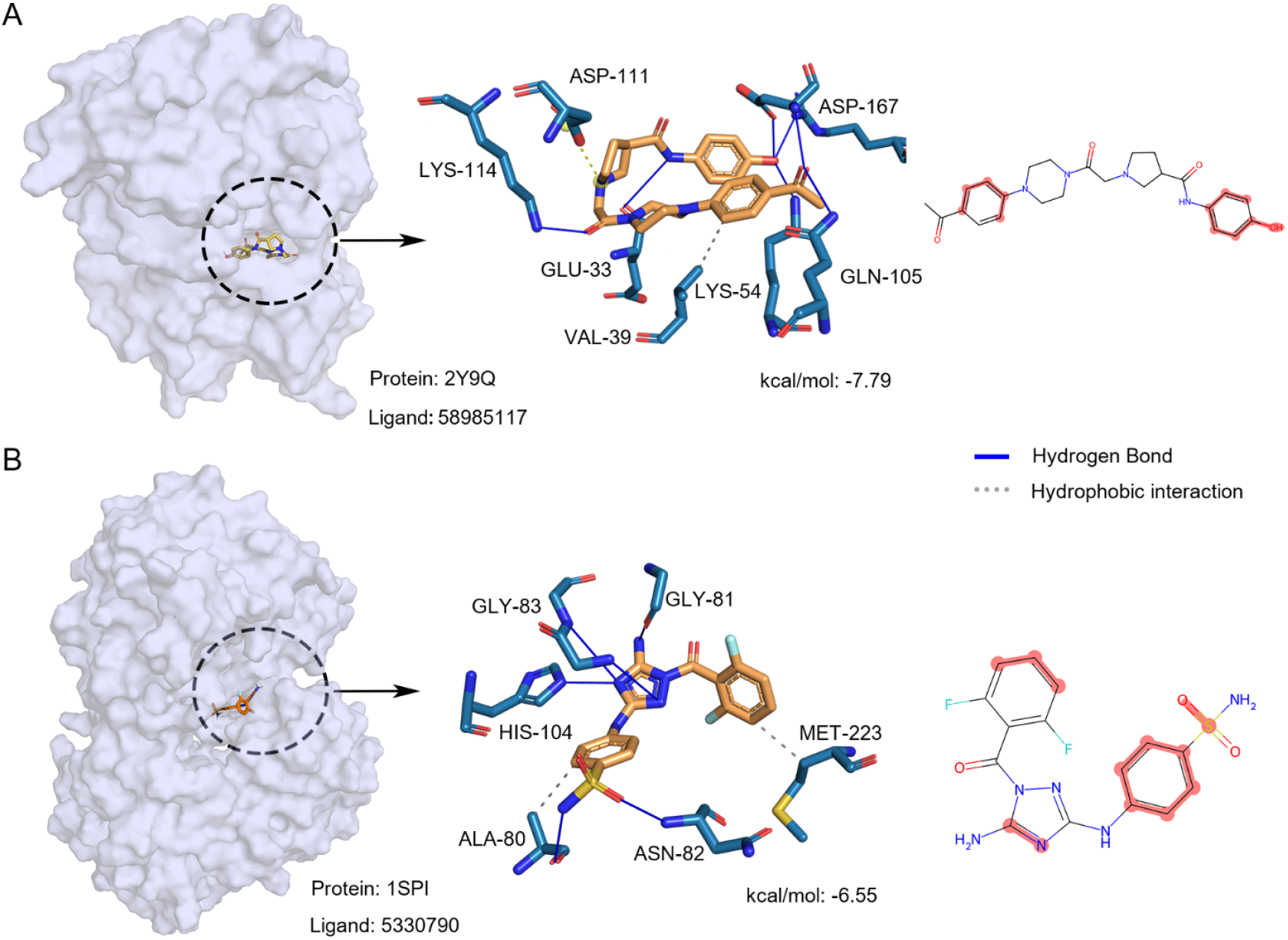
Molecular docking analysis of two examples. A. Docking of the protein Mitogen-activated protein kinase 1 to ligand. B. Docking of the protein Dual specificity mitogen-activated protein kinase kinase 2 to ligand.

### 3.6. Webserver and software development

To facilitate easy access to Kinhibit, we developed and published a user-friendly web server http://kinhibit.biotools.bio. The web server runs on a Linux server with 8 cores, 64GB memory, and 100GB hard disk. When submitting the ligand-kinase binding affinity prediction task, the user only needs to upload or paste the SMILES of the ligand. After the background calculation of the server is completed, the predicted ligand-kinase binding affinity for each input SMILES will be displayed in a tabular format. Users can download forecast results from the server in different file formats (EXCEL, PDF, and CSV). The “Help” page provides the usage guide of the web server for the convenience of users. Considering that the web server can run stably, our server has a limit on the number of SMILES submitted each time. If users need high-throughput predictions on large data sets, they can download a standalone version of the model on the download page for local predictions.

## 4. Conclusion

In this study, we proposed a novel framework combining graph contrastive learning and structure-informed protein language model (ESM-S) to improve the accuracy of kinase inhibitor binding affinity prediction. By jointly learning the features of ligands and kinases, this method effectively improves the characterization ability of ligands and verifies their superior performance on multiple MAPK signaling pathway kinases (RAF, MEK, and ERK). Experimental results show that our model outperforms existing computational tools, including BatchDTA, KIPP, AutoDock Vina, and GPT4Kinase. In addition, through interpretable analysis of the model, we found that this method is able to learn information from SMILES that helps to accurately predict ligand-kinase binding affinity. To facilitate the research of kinase inhibitors, we have also developed a user-friendly web technology platform. In general, our method provides support for bioinformatics research such as kinase inhibitor screening and drug discovery, and shows broad application prospects.

