## Supplementary material for "Kinase-Inhibitor Binding Affinity Prediction with Pretrained Graph Encoder and Language Model"

### Supplemental Materials

**Table 1.** Two label conversion strategies.

| **Strategy** | **Condition** | **Affinity** | **Label** |
|---|---|---|---|
| Strategy-A | 10,000 $\leq$ Kd | low | 0 |
| | 100$\leq$ Kd $<$10,000 | medium | 1 |
| | Kd $<$100 | high | 2 |
| Strategy-B | 5000$\leq$ Kd | low | 0 |
| | 100$\leq$ Kd$<$5000 | medium | 1 |
| | Kd$<$ 100 | high | 2 |

#### Performance evaluation

To fairly evaluate the prediction performance of the model, we refer to the ref.x(Liu et al., 2024), and apply the Area Under the Curve (AUC) and the following evaluation metrics:

$$Accuracy = \frac{TP + TN}{TP + TN + FN + FP} \tag{1}$$

$$Recall = \frac{TP}{TP + FN} \tag{2}$$

$$Precision = \frac{TP}{TP + FP} \tag{3}$$

$$\text{F1 Score} = \frac{2 \times TP}{2 \times TP + FP + FN} \tag{4}$$

Where $TP$, $TN$, $FP$, and $FN$ represent the number of true positives, true negatives, false positives, and false negatives, respectively.
